## Supplemental Table 1 for "Structure and function of the ROR2 cysteine-rich domain in vertebrate noncanonical WNT5A signaling"

**Supplementary Table 1. Data collection and refinement statistics for ROR2 CRD-Kr**

|  | **ROR2 CRD-Kr**  **(Native)** | **ROR2 CRD-Kr**  **(Pt-SAD)** | **ROR2 CRD-Kr**  **(S-SAD)** |
| --- | --- | --- | --- |
| **Data collection** |  |  |  |
| Beamline | DLS-I03 | DLS-I03 | DLS-I23 |
| Space group | P3_2_21 | P3_2_21 | P3_2_21 |
| Unit-cell parameters  a, b, c (Å)  *α, β,* *γ* (^o^) | 113.6, 113.6, 45.2  90.0, 90.0, 120.0 | 106.1, 106.1, 42.2  90.0, 90.0, 120.0 | 113.6, 113.6, 45.1  90.0, 90.0, 120.0 |
| No. of crystals / data sets | 1/1 | 2/2 | 1/1 |
| Resolution limits (Å - STARANISO) | a, b= 3.1, c = 2.3 |  |  |
| Wavelength (Å) | 0.97625 | 1.0500 | 1.7711 |
| Resolution (Å) | 94.93-2.43 (2.70-2.43) | 53.00-3.00 (3.10-3.00) | 56.80-2.95 (3.03-2.95) |
| No. of unique reflections | 7611 (381) | 5599 (397) | 7171 (519) |
| Completeness (%) | 92.6 (60.3) | 99.9 (99.5) | 98.6 (96.5) |
| Multiplicity | 9.2 (6.1) | 28.1 (24.3) | 184.7 (38.7) |
| ⟨*I*/*σ*(I)⟩ | 14.9 (1.6) | 23.5 (6.4) | 25.0 (1.6) |
| R_merge_ (%) | 6.9 (82.9) | 14.7 (59.6) | 44.2 (>100) |
| R_pim_ (%) | 2.5 (35.5) | 3.0 (12.3) | 3.2 (33.5) |
| CC_1/2_ | 1.0 (0.81) | 1.0 (0.8) | 1.0 (0.9) |
| **Refinement** |  |  |  |
| No. reflections (test set) | 8688 (430) | - |  |
| R_work_/R_free_ | 24.2 /25.6 | - |  |
| No. atoms |  |  |  |
| Protein | 1670 | - |  |
| Ligand | 10 | - |  |
| Mean B factor (Å^2^) |  |  |  |
| Protein | 112.0 | - |  |
| Ligand | 137.1 | - |  |
| RMSD bond lengths (Å) | 0.008 | - |  |
| RMSD bond angles (Å) | 0.91 | - |  |
| Ramachandran plot (%) |  |  |  |
| Favoured | 96.1 | - |  |
| Allowed | 3.4 | - |  |
| Outliers | 0.5 | - |  |

Data in parenthesis refer to highest resolution shell unless otherwise stated. RMSD: Root Mean Square Deviation. In the case of native data, ‘completeness’ to the ellipsoidally-truncated value from autoPROC+STARANISO.
