## Supplemental Table 2 for "Structure and function of the ROR2 cysteine-rich domain in vertebrate noncanonical WNT5A signaling"

**Supplementary Table 2**

| **Protein** | **Fz8-PAM**  **(4F0A)**^A^ | **Smo (5L7D)** | **Fz8**  **(1IJY)** | **sFRP3 (1IJX)** | **MuSK (3HKL)** | **NPC1**  **(3GKI)** | **RFBP (not in PDB)** | **FRα (4LRH)** | **JUNO (5EJN)** | **FRβ (4KMZ)** |
| --- | --- | --- | --- | --- | --- | --- | --- | --- | --- | --- |
| **ROR2** | 1.19^B^  81^C^  39.62^D^ | 1.30  78  35.87 | 1.30  83  37.19 | 1.28  83  36.53 | 0.88  110  62.08 | 1.94  74  27.99 | 2.40  58  17.20 | 2.50  54  15.39 | 2.43  50  15.10 | 2.38  51  15.57 |
| **Fz8-PAM (4F0A)** |  | 1.00  83  43.64 | 0.16  117  102.54 | 0.42  106  79.01 | 1.30  78  38.33 | 2.17  61  21.36 | 2.54  40  13.35 | 2.76  44  12.16 | 2.51  44  13.31 | 2.44  43  14.14 |
| **Smo (5L7D)** |  |  | 1.09  84  43.18 | 1.04  80  44.60 | 1.26  78  40.84 | 2.32  54  19.01 | 2.68  46  11.24 | 2.62  46  13.73 | 2.64  44  12.47 | 2.49  48  15.65 |
| **Fz8 (1IJY)** |  |  |  | 0.45  108  78.07 | 1.35  76  36.13 | 2.14  62  22.27 | 2.50  45  15.00 | 2.66  43  13.41 | 2.38  45  15.40 | 2.53  43  14.34 |
| **sFRP3 (1IJX)** |  |  |  |  | 1.36  79  36.65 | 2.41  60  17.28 | 2.31  45  17.60 | 2.59  47  14.13 | 2.49  39  13.87 | 2.47  46  15.62 |
| **MuSK (3HKL)** |  |  |  |  |  | 2.14  67  22.48 | 2.25  55  20.03 | 2.30  55  15.14 | 2.16  53  19.21 | 2.13  53  23.24 |
| **NPC1 (3GKI)** |  |  |  |  |  |  | 1.73  82  35.46 | 1.95  89  31.57 | 1.74  81  36.37 | 1.74  90  30.58 |
| **RFBP (not in PDB)** |  |  |  |  |  |  |  | 0.68  160  104.06 | 0.85  138  80.30 | 0.66  157  105.39 |
| **FRα (4LRH)** |  |  |  |  |  |  |  |  | 0.42  161  125.21 | 0.16  194  176.05 |
| **JUNO (5EJN)** |  |  |  |  |  |  |  |  |  | 0.46  157  120.83 |
